# Using a betting game to reveal the rich nature of visual working memories

**DOI:** 10.1101/2020.10.28.357442

**Authors:** Syaheed B. Jabar, Kartik K. Sreenivasan, Stergiani Lentzou, Anish Kanabar, Timothy F. Brady, Daryl Fougnie

## Abstract

When we ask people to hold a color in working memory, what do they store? Do they remember colors as point estimates (e.g. a particular shade of red) or are memory representations richer, such as uncertainty distributions over feature space? We developed a novel paradigm (a betting game) to measure the nature of working memory representations. Participants were shown a set of colored circles and, after a brief memory delay, asked about one of the objects. Rather than reporting a single color, participants placed multiple bets to create distributions in color space. The dispersion of bets was correlated with performance, indicating that participants’ internal uncertainty guided bet placement. Furthermore, relative to the first response, memory performance improved when averaging across multiple bets, showing that memories contain more information than can be conveyed in a single response. Finally, information about the item in memory was present in subsequent responses even when the first response would generally be classified as a guess or report of an incorrect item, suggesting that such failures are not all-or-none. Thus, memory representations are more than noisy point estimates; they are surprisingly rich and probabilistic.

## Introduction

Working memory refers to the capacity to keep information active and accessible when it is no longer present in the environment. This capacity is involved in nearly all domains of cognition and its central importance is reflected by its high correlation with important measures such as fluid intelligence^1,2^ and academic success^3,4^. Despite the unquestionable importance of working memory in our daily lives, it has a surprisingly limited capacity. For example, we struggle to remember simple features from a handful of visual objects^5,6^. Importantly, memory for simple features is correlated with memory for complex objects ^7,8^, enabling insights into everyday limitations in working memory through the study of memory for simple visual features, such as color, which can be presented in a well-controlled manner.

Contemporary theories have constructed elaborate models of visual working memory based on data from continuous-color adjustment tasks, which require participants to report the color of a remembered object from a color wheel or bar. When participants are shown just a few colors to memorize, responses are not veridical, but tend to be close to the correct value. However, with as few as five or six colors the reports often differ greatly from the presented color, suggesting that memory is limited ^9–12^. Researchers collect many responses per participant in order to generate aggregate distributions, which are often used to differentiate between models of working memory. While this approach has yielded many important insights on memory ^9, 13–16^, it precludes exploration of the nature of individual memories on individual trials — which is in many ways the core issue at stake in theories of memory representation.

For instance, one core issue is the complexity of memory representation of items. Are the noisy responses produced in adjustment tasks the result of noisy point estimates, or are memory representations much richer? Some argue that most of the sensory representation information is lost by the time a decision has to be made^17^, such that memories consist of a point representation only (i.e., I saw red, but now all I know is that I think it is pink) or a point estimate that may be associated with a non-probabilistic notion of confidence. Models of visual working memory where memories ‘drift’ over time^18^ or with noise-less representations^19^ often assume this is true. However, another possibility is that memory representations are richer and more complex than simple point representations, consisting, for example, of stored probability distributions over a feature space^10, 20–22^. If this is true, and if such richness is accessible to participants’ decisions, then it is incomplete and potentially misleading to assume that the discrete response is equivalent to the memory. In particular, if responses are noisy samples ^23,24^ from a probability distribution, then existing estimates of memory constraints could be significantly underestimating the amount of information represented in memory. Some models predict such complexity: for example, population-based models can have rich potentially asymmetric internal distributions in feature space, at least in displays with multiple items being represented simultaneously^22,25,26^. In such cases, there is no straightforward conversion of the probability distribution to a single response.

In order to explore the degree to which individual representations are rich and complex (akin to the across-trial aggregate distributions observed in adjustment tasks) it is necessary to get more information from a participant than a single estimate. We developed a betting game task (Figure 1) that encourages participants to report the properties of the uncertainty over feature space of individual memory representations (*uncertainty profiles*). Specifically, participants were given a working memory task of five colored circles and then asked to create a distribution over the color space of the potential color of a randomly probed item. While this uncertainty profile was created from six discrete responses (placing six discrete circular Gaussians), there was immense freedom in the final distribution that participants could draw via this method. Essentially, participants were encouraged to generate a distribution that was proximate to the internal distribution of uncertainty for that memory (assuming such a distribution is accessible to the participant). Participants were rewarded by the height of the distribution over the correct value, reinforcing participants to construct something akin to an accurate probability distribution.

**Figure 1.**
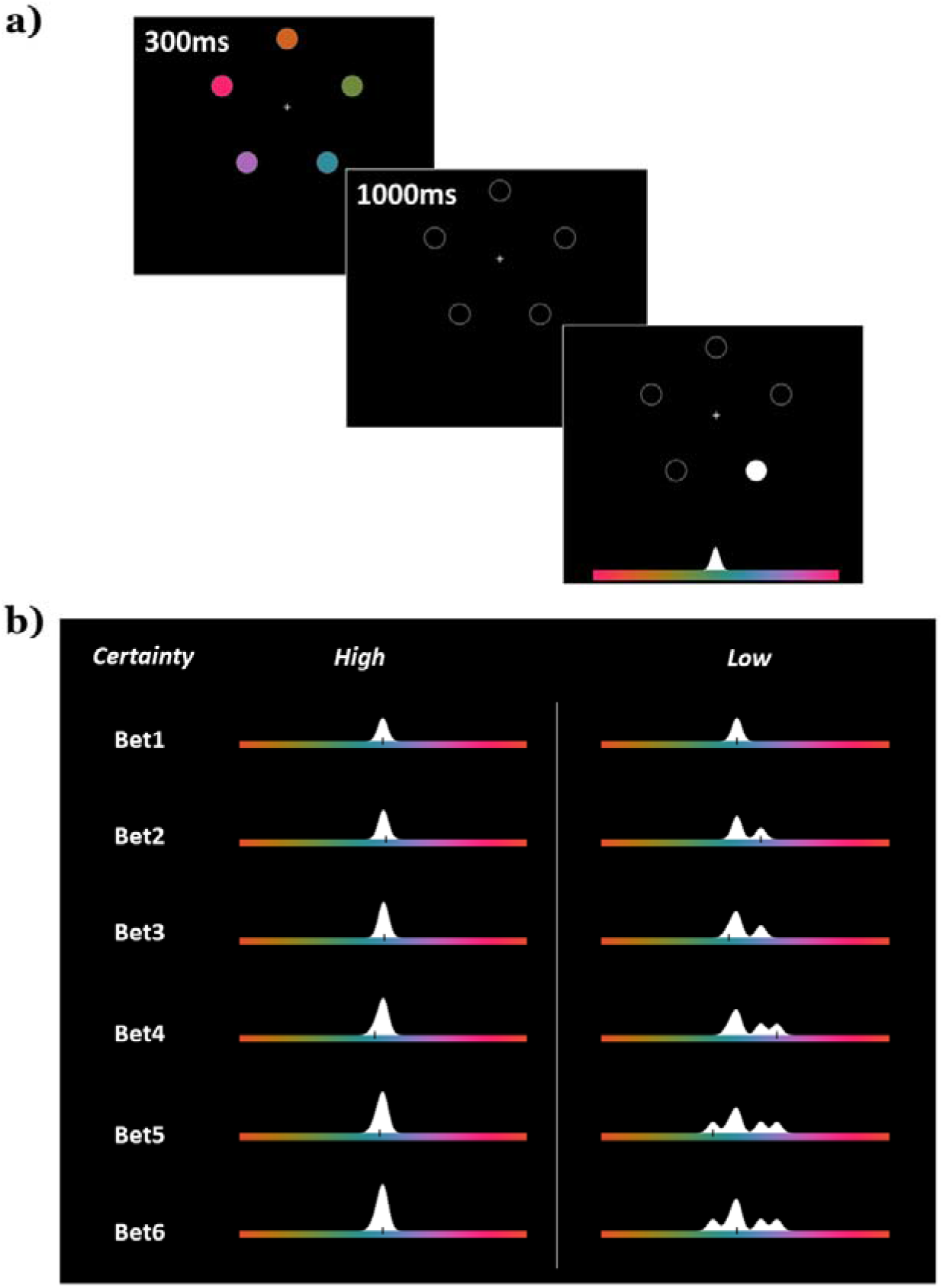
Experiment protocol. a) Five color targets were shown, followed by a delay period. A color response bar then appeared, and one of the target locations highlighted. In Experiment 1 participants were always asked to make 6 ‘bets’, b) In the first bet, participants used a mouse to laterally shift the colors of the response bar. Participants were asked to place the color they remembered in the cued location at the center of the bar, where the white Gaussian is. Participants confirmed their choice with a mouse click. For the 2^nd^ to 6^th^ bets, another Gaussian (with half the height of the original), appeared at a random position on the bar, and participants were asked to move the mouse to place the Gaussian over what they thought was the color they remembered, again confirming the bet with a mouse click. This was repeated for six bets per trial. Note how participants were free to spread out the bets as they wished. How the profile would look like if they confirmed the bets was always previewed to participants. If they were very certain, they could stack the bets, forming a narrow uncertainty profile (left column). Doing this could earn them more points if they were on-target, but at the risk of no reward if there was no height of the drawing over the target color. In this case, participants would have received 132 points. If uncertain (right column), participants could spread the bet, as a lower-risk, but lower-reward option (in this case receiving 21 points due to the reduced height at the target color).

Using this new task, we found that participants created uncertainty profiles that were close facsimiles to the error distribution across trials. Moreover, the placement of bets tracked internal uncertainty—when bets were more widely spaced, participants were less accurate, and *vice versa.* This suggests that participants have some access to the underlying uncertainty in a memory representation and could utilize this information when creating the uncertainty profiles. Most provocatively, we found evidence that participants had more information about the memory items than was contained in the first report: representations derived across multiple responses were significantly closer to the true stimulus color than the first response alone, and trials classified as errant responses based on participants’ first report (either guess or swap responses [e.g., where they reported yellow for a blue item]) still contained information about the true stimulus color in their uncertainty profile. Taken as a whole, these findings reveal that participants do not have access to only a discrete value in memory, but instead a complex probability distribution. Moreover, the results suggest that existing measures of working memory capacity and performance are underestimating the amount of information that we retain.

## Results

### Evidence that the reported distributions incorporate uncertainty

To explore the degree to which participants’ bets captured their memory uncertainty, we first looked at whether the spread of bets predicted memory performance. This analysis leverages the findings that memories vary in quality^13,14^ and that participants have some metaknowledge of the uncertainty in memory ^27–30^. Thus, to maximize performance, a participant should place bets narrowly when the uncertainty of the true color is low and spread bets widely when uncertainty is high. Indeed, we found significant positive correlations (*p* <.05) between the magnitude of the error of the first response and the median absolute distance between adjacent bets for 32 of the 34 participants (mean *r* = .393). These individual correlations were Fisher-transformed (mean *z* = .430, 95% *CI* = [0.010, 0.977]) and a *t*-test found this distribution to be significantly different from zero (*t*(33) = 12.57, *p* <.001). Similar results were found using other measures of the bet spread, including the standard deviation (mean *z* = .379) or the interquartile range ^31^ (mean *z =* .362) of the uncertainty profile.

This analysis demonstrates that the way the bets are placed reflects the error in participants’ first response. But how closely does the reported uncertainty within trials match participants’ across-trial error, as assessed by the profile of the error in first responses across trials? If participants were accurately recreating the uncertainty in memory and sampling from this to derive first responses^23^, the across trial error distributions would match the average of the uncertainty profiles reported within a single trial. We examined this using two-sample Kolmogorov-Smirnov (*KS)* tests ^32^ to compare the across-trial error distribution (error of the first response relative to the correct answer) to the trial averaged uncertainty profile for each participant. Uncertainty profiles were circle-shifted to align the target colors and averaged at each integer value of the color space (e.g., at 360 discrete points). For 30 of the 34 participants, the *D*-value (the *KS* statistic which measures the maximum difference between the two empirical cumulative distribution functions) was small (mean *D* = .041, *SD* = .015) and non-significant (p>.05), highlighting that the average reported uncertainty was not significantly different from the across-trial error of the first response (Figure 2). As a bootstrap control, surrogate *D* values for the null hypothesis were simulated by drawing two sets of 1000 samples from the average of the two distributions per participant. *KS* tests on these theoretically identical sets should indicate to us the expected range of *D* values under the null hypothesis. We carried out a thousand of these simulations and found that the actual *D* value fell within the range of control *D* values (95% confidence interval = [019, .064]). In other words, the two distributions (the trial averaged uncertainty profile and the first bet error distributions) did not significantly differ in shape.

**Figure 2.**
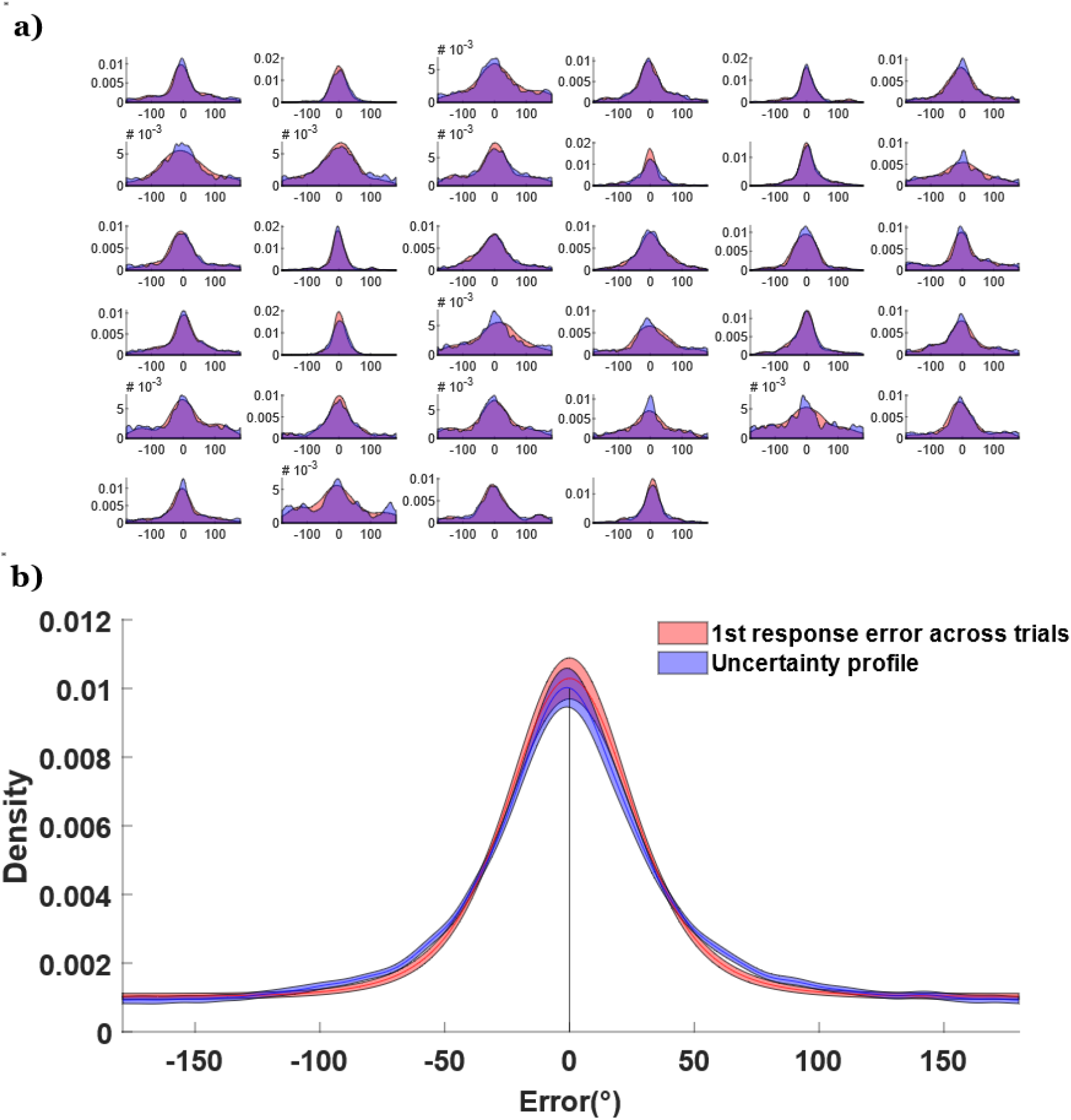
Comparing the error distribution of the first response across trials (red) versus the uncertainty profiles (after all six bets) averaged across trials (blue). a) The comparison between the two distributions for each of the 34 participants. b) Aggregate of the distributions across participants. The shaded region represents 1 standard error above and below the mean.

### Evidence that the bets contain more information than found in the first response

The previous analyses highlight that the created distributions convey rich information about memory uncertainty. But do they also contain information about what is in memory beyond what is captured by the initial response? We analyzed the placement of individual bets, as well as the cumulative circular average of the bets (e.g. if the first bet was a color 10° clockwise and the second was 4° counter-clockwise, the cumulative average would be 3° clockwise). As shown in Figure 3, the first bet placement is the most accurate, with a relatively monotonic decline in the accuracy of individual bets across the trial (when considering the center of bets in isolation). To examine this, we subjected the bets to a one-way ANOVA with the six levels of bet order as a within-subjects factor. Errors generally increased with bet number (bet 1 = 31.0°, bet 6 = 32.5°), *F*(5,165) = 3.39, *p* =.006, *η*^2^ =.093. This result is not surprising given that the first response was worth double the points of subsequent responses and that subsequent responses occurred after increasing delays and potential interference from previous responses.

**Figure 3.**
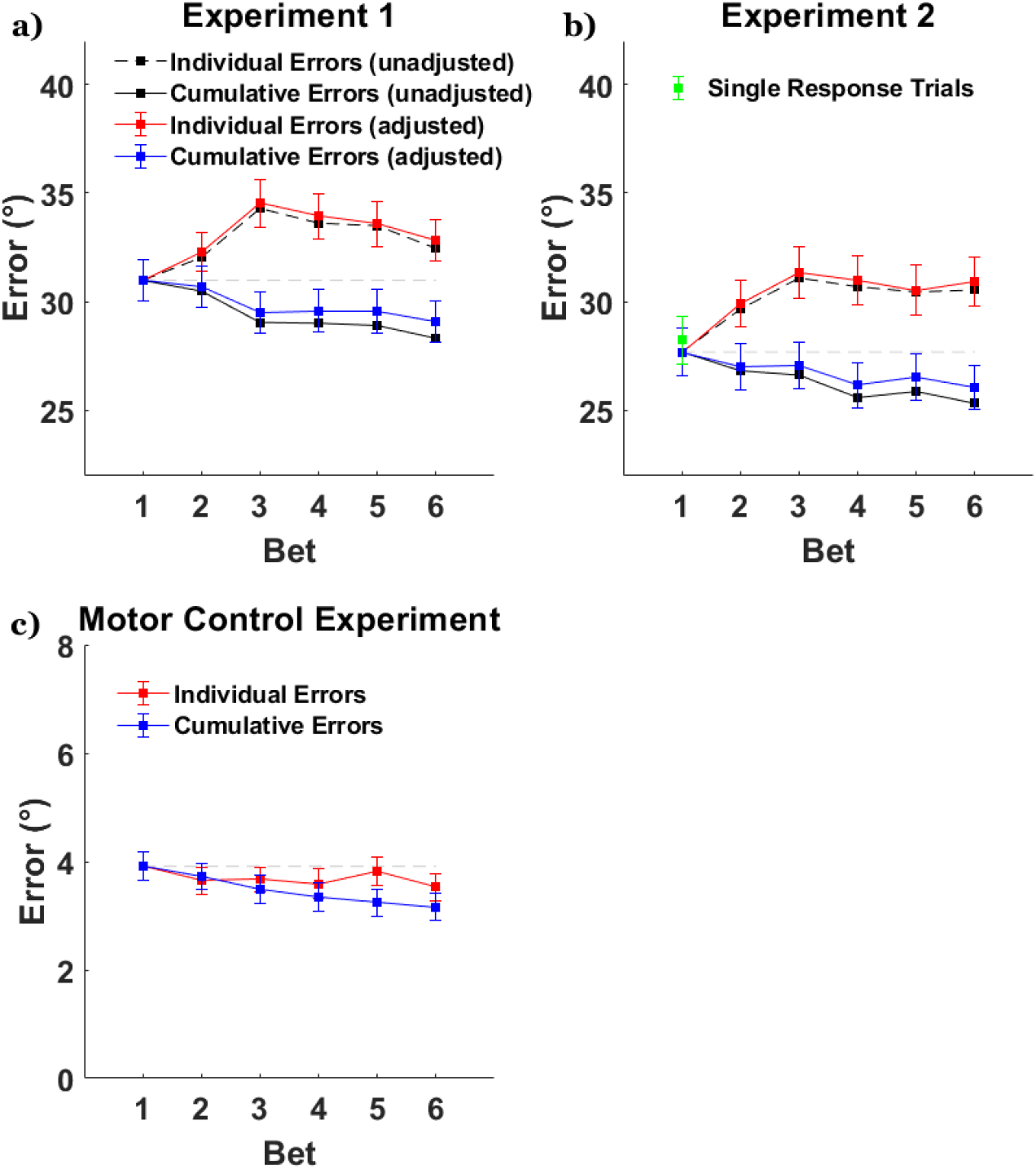
Individual response errors (red) and cumulative errors (blue) as a function of response order. Cumulative errors are calculated as the error of the mean of responses. For example, the cumulative error for response 3 would be influenced by response 2 and response 1, whereas the individual response error for response 3 is solely determined by that bet. a) Errors for Experiment 1. Adjustments were made based on the results of the motor control experiment to try to minimize the impact of motor noise. b) The equivalent errors for Experiment 2. The green marker is the mean for the single response trials. c) Errors for the motor control experiment where the stimuli stayed on-screen.

Despite being less accurate than the first responses, subsequent responses could nevertheless contain additional, unique information about the target item^24^. To test this, we examined whether additional bets were providing novel information not contained in the first response by testing whether the center of the uncertainty profile approaches the true color with repeated placement of bets. We subjected mean bet placement to a one-way ANOVA and found that performance improved with additional bets (bet 6 = 28.3°), *F*(5,165) = 5.35, *p* < .001, *η*^2^ =.140, despite the fact that average error trended worse for the later bets, suggesting that the combination of responses contains more information than any individual response, including the first response.

One possibility is that the benefit of averaging is solely due to averaging out individual noisy motor responses. In this view, participant’s full knowledge of the item is captured in the first response, but the response process is noisy, and this noise is cancelled out over the course of multiple responses. The idea that motor noise is present and would be averaged out by multiple responses is reasonable. However, response error is unlikely to be sufficiently large enough to play much of a role in effecting the large response error for remembering 5 items. To test this directly, we utilized the data from the motor control task. Unsurprisingly, we observed a benefit when combining multiple responses, *F*(5,80) = 15.6, *p* <.001, *η*^2^ =.0.49, but the change in error was less than a degree (bet 1 vs all 6 combined: difference = 0.75°).

Importantly, we can subtract the observed benefit from this control study from that observed in Experiment 1. The mean error for each bet-value obtained from this control experiment was removed from each Experiment 1 participant’s error. Hypothetically, if a participant had a mean cumulative bet error of 36° at bet 1 and 33° when considering all 6 bets, then removing the mean bet 1 (3.91°) and cumulative benefit from all 6 responses in the motor control experiment (3.16°) would suggest that there is still approximately a 2.25° decrease in cumulative error that cannot be attributed to averaging across motor noise. This adjustment was done for all participants with both the cumulative and the individual bet errors (the black vs colored lines in Figure 3a show the effect of adjusting the means based on motor error). Having factored out motor error, we still found evidence of reduced error when averaging bets, *F*(5,165) = 2.86, *p* =.017, *η*^2^ =.080. This suggests that there is additional information in the bets not captured by a single response, and while a benefit is expected when simply averaging noisy responses, it is insufficient to fully explain the effect seen in Experiment 1.

A related concern is that participants are spending insufficient effort on the first response because they can “make-up” for an inaccurate first response in subsequent responses. Several pieces of evidence argue against this. First, performance for our first response is expected given previous results in the literature. This is most clearly seen when we decompose our errors using mixture models to facilitate comparisons to previous work (using MemToolBox ≡3). We find a guess rate of 37.1% with a set size of five, which is between the 16% found with set size 3 and 59% found with set size 6 in Zhang & Luck^12^. The amount of guessing reflects a capacity of 3.15 (compared to a capacity of 2.46 found in Zhang & Luck^12^). Precision *SD* in our online task after factoring out ‘guesses’ was approximately 29.4°, which is also close to our previous (unreported) in-lab iteration of the betting task (with set size of 5, mean guess rate = 31%, *SD =* 28.1°). In addition, our first bet was weighted higher than others to encourage participants to emphasize this response. However, to address this issue more fully we ran a replication study that interspersed single- and multi-bet trials (intermixed).

### Effect of interspersing single response trials

We replicated the results under conditions in which participants were uncertain if they were to make one report or multiple bets (Experiment 2). The first bets in Experiment 2 (*M* = 27.6°, *SD* = 12.0°) did not significantly differ in error magnitudes compared to Experiment 1 (*M* = 31.0°, *SD* = 11.3°), *t*(63) = 1.13, *p* = .260. The single response trial errors (*M* = 28.2°, *SD* = 12.0°) did not differ from the errors on the first bet of the multiple bet trials in error magnitude, *t*(30) = 0.56, *p* = .582. For the individual errors of the multiple bet trials, errors increased from bet 1 to bet 6 (*M* = 30.5°, *SD* = 12.6°), with the one-way ANOVA showing *F*(5,150) = 3.45, *p* =.006. For the cumulative errors of the multiple bet trials, errors decreased from bet 1 to bet 6 (*M* = 25.3°, *SD* = 11.1°), with the oneway ANOVA showing *F*(5,150) = 5.47, *p* <.001, *η*^2^ =.154. Adjusting these cumulative errors by the motor benefits still suggested a decrease, *F*(5,150) = 2.63, *p* =.026, *η*^2^ =.081.

The findings replicate the benefit of averaging bets and provide additional evidence against the possibility that the observed decrease in cumulative error as a function of bet was due to participants underperforming during the first report.

### An argument for uncertainty profiles: The case of ‘guess’ and ‘swap’ errors

Thus far we have focused on how the uncertainty profiles provided by participants generally match their aggregate performance, with the cumulative error analyses further suggesting that these profiles contain more information than any single response. However, the benefit of measuring these profiles is that it affords the ability to explore novel issues, like how much information is available about the target when participants’ initial report is far from the target.

Many models of visual working memory make claims that distinct ‘states’ underlie some kinds of memory errors. For example, some models suggest that far away responses reflect ‘guesses’, where no target-specific information is available to participants^12^. Other models suggest that some far away reports reflect ‘swaps’, where participants report a non-target item ^9,34^, either erroneously^34,35^ or strategically^36^. The betting game task allows us to examine whether these responses truly reflect distinct states with no target-specific information by more closely examining the uncertainty profiles for such “swap” trials and “guess” trials.

We classified trials into three trial types: target, swap, and guess trials^34^. Unsurprisingly, based on the first bet errors the majority of trials were classified as target trials (*M* = 73.3%). There were significantly more guess trials (*M* = 20.1%) than swap trials (*M* = 6.6%), with the target trials being the majority (all comparisons, *p*<.001). The lack of swap errors likely stemmed from the fact that the locations of the colors were fixed and predictable across trials ^9,35^. Only 33 of the 65 available data sets had trials that could be classified as swap trials. In order to have sufficient power to examine swaps, we combined the data from Experiment 1 and the multiple bet trials from Experiment 2. Swap trials (*M* = 39.1°) did not significantly differ from guess trials (*M* = 43.8°) in terms of bet spread, *t*(26) = 0.88, *p* = .387. When combined with our previous finding that bet spread was correlated with initial response error and thus may reflect memory uncertainty, this result is consistent with other work ^36^ demonstrating that confidence ratings do not differ between swap and guess trials. In contrast, the bet spread for the target trials (*M* = 24.2°) was significantly smaller than the two other trial types (*p*s < .001). Most importantly, one-way ANOVAs revealed that *all three* trial types had a decrease in cumulative bet errors from bet 1 to bet 6 (*p*s < .001, *η*^2^s > .01). This suggests that single response methods tend to underestimate the amount of information about the target item that participants store not only for on-target reports, but also for swap or guess reports -- and that participants do indeed have target-specific information on such trials.

Residual target encoding in cases of ‘swap’ and ‘guess’ errors can be further examined via participants’ bet placement over the target color. For example, if a ‘swap’ reflects a complete replacement of the tested item (at either encoding, storage, or retrieval), then we expect the height of the uncertainty profile over the target color to be no higher than over other random positions. To examine this, we took the uncertainty profiles for each trial and centered them on the first response such that errors biased towards the target side were coded as positive. We found that the sum of heights over the estimates biased towards the direction of the target were higher than estimates biased away, for all three target types (Figure 4a, all *p*s < .001, *η*^2^s > .31). Finally, we compared the height of the distribution over the target to the height over a “control” color that was equidistant from the first bet but in the opposite direction as the target (Figure 4b). If the profiles contain no target information, these two heights should be identical. Instead, bet height was significantly greater for target versus control color on guess and target trials (both *p*s<.001, *η*^2^s > .23) and marginally significant for swap trials *t*(32) = 1.80. *p* = .081, *η*^2^ = .09 (notably, swap trials had reduced power due to low trial counts).

**Figure 4.**
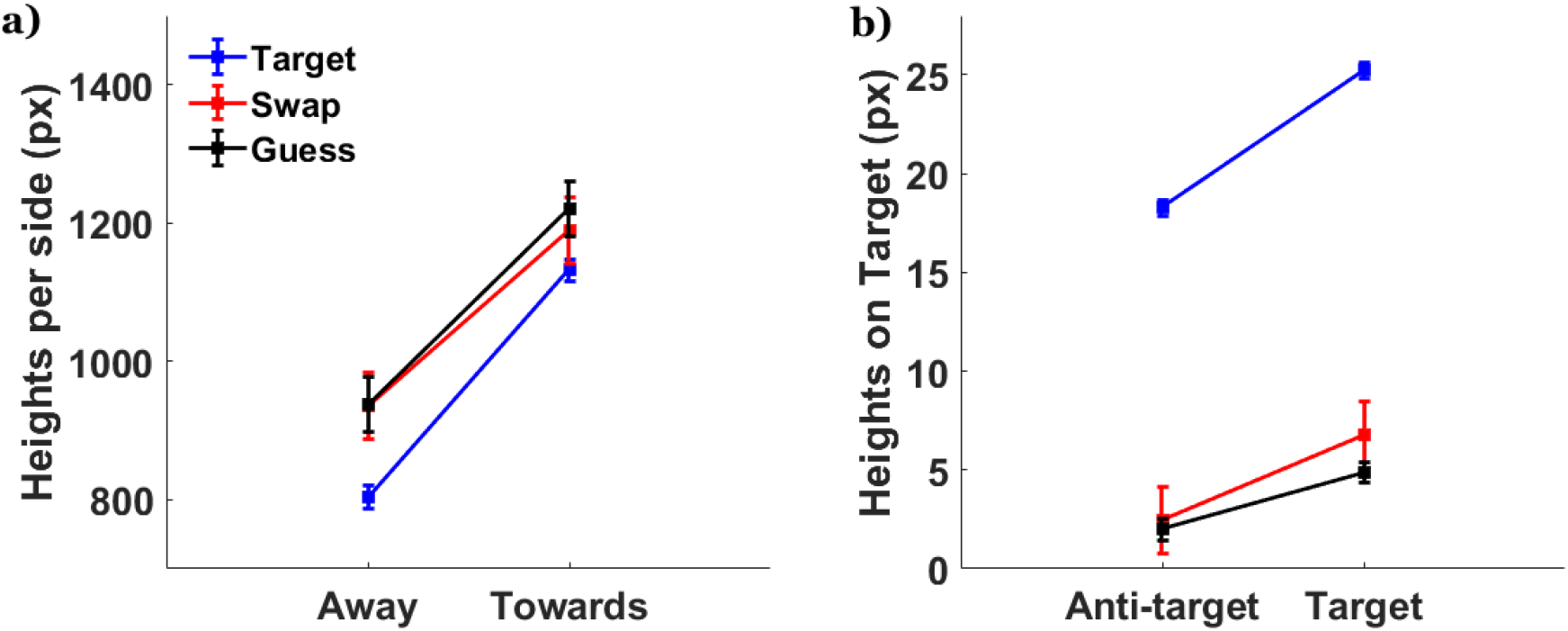
Post-classification analyses. a) Sum of profile heights either towards or away from the target. b) Height over the actual target compared to the anti-target (the color that is equidistant to the target from response 1, but in the opposite direction).

We find that participants encode and report something meaningful about target identity, even when their first response performance would suggest otherwise. Thus, relying on single responses can cause an underestimation of the richness of mental representations: target encoding can be readily observed even in trials that would traditionally be classified as ‘swap’ or as ‘guesses’.

## General Discussion

There has been considerable focus on understanding the nature of visual working memory. Most of this work has focused on categorizing the structural limitations of our working memory system— How much information are we able to store and what is the unit of this limit (e.g., objects, features, bits of information)? However, these questions have been investigated with analysis techniques that examine the average memory state across many trials, not individual memories; behavioral estimates of memory performance rely exclusively on tasks that cannot differentiate whether participants are encoding a discrete value (e.g., a shade of red) or something more complex and probabilistic. An underlying assumption of many models, particularly physiologically plausible models, is that representations consist of patterns of activity that can be effectively described as probability distributions that exist over some feature space^22,25,26^. Given this, it is unfortunate that there has been a near exclusive focus on evaluating and comparing theoretical models using tasks in which single, discrete responses or estimates are the sole basis of performance. At best, these discrete responses are capturing a representative summary of this probability distribution (e.g., the mode of a Gaussian). However, this optimistic scenario might not be accurate. Discrete reports may represent something besides the best estimate of the memory, or the underlying distributions may be sufficiently complex that any method of condensing them into a single response throws away considerable information.

To explore this, we developed a betting game task where participants were asked to do more than render a single discrete judgment. Instead, participants placed six independent Gaussian bets over a color-feature space, which were combined cumulatively to form a final response distribution that was rewarded. The trial-specific distributions drawn revealed that participants could access information about item uncertainty on a trial-by-trial basis. We found considerably larger response error for trials in which bets were widely spaced (indicating high uncertainty) than for trials that were closely spaced (indicating low uncertainty). Moreover, the uncertainty profiles drawn by participants on individual trials were comparable to the across trial aggregate distributions, suggesting that participants are tapping into something akin to an internal probability distribution over color space to guide bet placement. This finding is in line with previous studies showing that participants have metaknowledge of memory quality ^27–30^ but goes further in also suggesting that this uncertainty distribution can be directly measured in a task with a betting game.

The uncertainty distributions revealed more than just trial-by-trial memory uncertainty. The distributions contained more information about the memory item than was captured in a single response. Specifically, the average response error for the mean of the uncertainty distribution decreased monotonically from the first to the last bet, despite the fact that the average error of each bet increased. This suggests that there was additional information contained in subsequent bets not contained in the first response. This was true even when participants did not know if they were placing one or multiple responses, and even after accounting for any contribution of motor error at response. Furthermore, even in trials that would have traditionally been classified as ‘swap’ or ‘guess’ trials based on a single response^9,12^, participants demonstrate that they have information about the target. The present findings highlight that existing measures and methods to estimate information storage capacity are problematic and may underestimate the contents of memory. An assumption in behavioral studies of working memory is that the responses gathered in memory tasks are a veridical reflection of the contents of memory. However, our data suggests that memory representations are richer than is captured by a single response.

All manner of working memory tasks require decision-making. A common strategy for studying working memory has been to minimize these decision-components and to treat the behavioral output of tasks as synonymous with the internal representation. We advocate a different approach. Given that working memory representations are rich and complex, it is impossible to derive a task where the output can be assumed to be a pure reflection of the contents of memory. Instead, researchers must consider how internal representations are converted into discrete responses and use tasks where such transformations can be tested empirically. Rather than limiting or complicating the study of working memory, this line of inquiry may offer a novel way to differentiate amongst theoretical models^27^.

Our findings by no means completely answer the myriad of questions about the capacity and nature of working memory. Instead, the present work highlights a novel framework and suggests new questions that we should be turning our attention towards. Theoretical or computational models with sufficiently specified mechanisms ought to predict what an internal distribution of uncertainty (on a trial-by-trial basis) might look like. For example, many models propose rich representations composed of uncertainty distributions over feature space ^22,37–39^, such as how competing encoded representations might interfere or influence one another in biologically plausible ways^26,35,40^. For example, models where population coding is taken as the basis of memory -- perhaps in early visual cortex for simple visual working memory tasks ^41–42^ -- memory representations may best be thought of as entire distributions across feature space, reflecting the activity of the entire neural populations^26^. Instead of simply acquiring single response data, the present technique (or something similar) could provide a novel means of testing such theories.

Research that takes seriously the question of what an individual memory looks like will help to bridge the gap between cognitive models on the one hand and biologically inspired neural models on the other. Given that our memories are rich and complex, our report methods would do well to embrace this complexity.

## Methods

### Experiment 1

#### Participants

Forty naïve participants (26 male, median age 23.5) were recruited for Experiment 1. All participants declared normal or corrected-to-normal visual acuity and color vision. Participants were recruited online (via www.prolific.co) for a base allowance of 5.20GBP per hour, and were told that they could also receive a monetary bonus depending on their performance (mean bonus = 2.00 GBP). The experiments were approved by the New York University Abu Dhabi Institutional Review Board. While similar in-lab pilot studies have shown the effects of interest with about 20 participants, we collected data from twice the number of participants to account for the possibility that online testing would increase noise in the data. As six participants had abnormally large error magnitudes (>66°) and were randomly placing bets, we removed them from the analyses.

#### Apparatus and stimuli

Because of the online nature of the experiment, stimulus sizes would vary slightly depending on the participant’s screen size/viewing distance. Assuming that a 27-inch 1920 × 1080 resolution monitor placed 70 cm away was used, targets would be located approximately 4° away from the center of the screen, with the target circles themselves occupying 1° of visual angle. The background of the screen was black throughout the experiment. Targets appeared in the same five equidistant locations (see Figure 1a for positions). The response color bar was approximately 6° from the center of the screen with a thickness of 1.0°. Effort was made particularly to ensure that the drawn display fit within the participant’s browser page. Participants were also asked to maximize their browser window prior to launching the experiment. The colors chosen for a given trial and for the response bar were sampled from the MemToolbox^33^ 360° color space, translated into RGB assuming an equal energy whitepoint. All reported experiments were programmed entirely in HTML Canvas/Javascript.

#### Procedure

In Experiment 1, each trial began with a blank screen with a fixation cross lasting for 500ms. Five colored stimuli then appeared on-screen for 300ms. These five colors were randomly chosen for each trial. Stimulus presentation was followed by another blank screen for 1000ms, during which time participants maintained the color information in working memory. After this delay period, the location of one of the five targets (each location equally probable) was then cued as the tested location. The response color bar appeared and for the first bet, participants used a mouse to laterally shift the colors of the response bar. Participants were asked to place the color they remembered to be in the cued location to be at the center of the bar, where a white distribution that is doubled the height of the standard Gaussian was located (standard deviation of 6° in color space). Participants confirmed their choice with a mouse click.

For the 2^nd^ to 6^th^ bets, a standard Gaussian appeared at a random position on the bar, and participants were asked to move the mouse to place the Gaussian over what they thought was the color they remembered, again confirming the bet with a mouse click (iteratively, for a total of six bets). Note participants were free to spread out the bets as they wished. During each bet, the updated uncertainty profile (based on current mouse position) was updated per frame and previewed to participants. Participants were instructed to build a distribution that matched the uncertainty of their internal state. The six Gaussians summed to create a final uncertainty profile for that trial. Participants were free to stack responses on top of one another (see Figure 4b, left column) or to spread it across a larger color space (right column). As a result, the uncertainty profile ‘drawn’ at the end of the sixth response need not have a Gaussian shape. Note that the Gaussians spawned in a random location between responses to prevent stereotyped successive clicks on the same color location, and participants had to wait a minimum of 300ms and make a lateral mouse movement before they could register the next click.

To encourage participants to report something resembling the true internal uncertainty, participants were awarded points based on the height of the final drawn distribution over the target color:

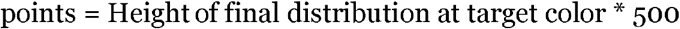

The *first* bet was twice as tall (i.e., worth twice as many points) to encourage participants to place the first bet accurately. To prevent participants from adopting the strategy of stacking responses at the peak of uncertainty, we implemented diminishing returns to stacking by reducing the value or rewarded points when stacking bets on existing bets. This penalty was scaled by height. Specifically, the height of the Gaussian component to be added was penalized by the current height of the drawn profile raised to a penalty parameter. For example, if the current bet is *b,* and the profile already drawn due to the previous bets is *y,* then the new bet *y’* is:

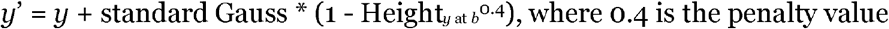

The taller the current height at the new bet value, the smaller the possible gain in height (and therefore the smaller the gain in points). Importantly, this penalty was built into the visualization of the drawn distribution seen by participants. After the six guesses were placed, the trial score, total score, as well as the current accumulated monetary bonus (every 200 points earned 0.10 GBP) were displayed on the feedback screen until the participant re-centered the mouse on the central fixation. The maximum possible points per trial was 146. Each participant first completed 6 practice trials, followed by 150 main trials, with a break every 50 trials. Practice trials were excluded from the analyses.

### Motor Control Experiment

A control study of Experiment 1 was done on 20 additional naïve participants (10 male, median age 23.5). We used the same betting protocol as Experiment 1 but stripped away the memory demands by leaving the stimulus on screen during the delay and report in order to isolate the error involved in reporting an onscreen stimulus. Because precision was expected to be high (therefore data low in noise), participants only were required to do 80 multiple response trials, preceded by 6 practice trials. Three participants were removed due to random betting, and the remaining participants had a very small magnitude of error (first bet mean absolute error = 3.91°).

### Experiment 2

As a replication of Experiment 1 and also to minimize the possibility that the predictability of always having to make multiple responses affected the precision of the first bet, Experiment 2 was conducted using 40 additional naïve participants (29 male, median age 23). We used the same betting protocol as Experiment 1 except that we randomly interspersed trials that required only a single response (i.e., just the first bet) amongst the trials that required placing a sequence of 6 bets. Participants completed 100 trials of each type. Critically, the response condition was not known until the second response; thus, participants could not anticipate at the time of the first response whether they would have the opportunity to make multiple bets. Of the 40 participants, 9 made random clicks (median absolute bet error > 66°, same cutoff as Experiment 1) and were dropped from the analysis.

## Acknowledgements

We would like to thank Karima Raafat for her comments on a previous version of the manuscript. This work was supported by NYUAD Research Enhancement Fund REF-175.

## Data Availability

The data that supports this study have been deposited at https://osf.io/7srv4/

## Code Availability

HTML/Javascript code for a demo version of the task is available at https://osf.io/7srv4/

## References

1. Fukuda, K., Vogel, E., Mayr, U., & Awh, E. (2010). Quantity, not quality: The relationship between fluid intelligence and working memory capacity. Psychonomic bulletin & review, 17(5), 673–679.

2. Kane, M. J., Bleckley, M. K., Conway, A. R. A., & Engle, R. W. (2001). A controlled-attention view of working-memory capacity. Journal of Experimental Psychology: General, 130(2), 169–183.

3. Alloway, T. P., & Alloway, R. G. (2010). Investigating the predictive roles of working memory and IQ in academic attainment. Journal of experimental child psychology, 106(1), 20–29.

4. Daneman, M., & Carpenter, P. A. (1980). Individual differences in working memory and reading. Journal of Memory and Language, 19(4), 450.

5. Luck, S. J., & Vogel, E. K. (1997). The capacity of visual working memory for features and conjunctions. Nature, 390(6657), 279–281.

6. Vogel, E. K., Woodman, G. F., & Luck, S. J. (2001). Storage of features, conjunctions, and objects in visual working memory. Journal of Experimental Psychology: Human Perception and Performance, 27(1), 92.

7. Palmer, J. (1990). Attentional limits on the perception and memory of visual information. Journal of Experimental Psychology: Human Perception and Performance, 16(2), 332.

8. Awh, E., Barton, B., & Vogel, E. K. (2007). Visual working memory represents a fixed number of items regardless of complexity. Psychological science, 18(7), 622–628.

9. Bays, P. M., Catalao, R. F., & Husain, M. (2009). The precision of visual working memory is set by allocation of a shared resource. Journal of vision, 9(10), 7–7.

10. Ma, W. J., & Jazayeri, M. (2014). Neural coding of uncertainty and probability. Annual Review of Neuroscience, 37, 205–220.

11. Mazyar, H., Van den Berg, R., & Ma, W. J. (2012). Does precision decrease with set size? Journal of Vision, 12(6), 10–10.

12. Zhang, W., & Luck, S. J. (2008). Discrete fixed-resolution representations in visual working memory. Nature, 453(7192), 233–235.

13. Fougnie, D., Suchow, J. W., & Alvarez, G. A. (2012). Variability in the quality of visual working memory. Nature communications, 3(1), 1–8.

14. Van den Berg, R., Shin, H., Chou, W. C., George, R., & Ma, W. J. (2012). Variability in encoding precision accounts for visual short-term memory limitations. Proceedings of the National Academy of Sciences, 109(22), 8780–8785.

15. Zhang, W., & Luck, S. J. (2009). Sudden death and gradual decay in visual working memory. Psychological Science, 20(4), 423–428.

16. Zhang, W., & Luck, S. J. (2011). The number and quality of representations in working memory. Psychological Science, 22(11), 1434–1441.

17. Yeon, J., & Rahnev, D. (2020). The suboptimality of perceptual decision making with multiple alternatives. Nature Communications 11, 3857.

18. Panichello, M. F., DePasquale, B., Pillow, J. W., & Buschman, T. J. (2019). Error-correcting dynamics in visual working memory. Nature Communications, 10(1), 1–11.

19. Cowan, N. (2001). The magical number 4 in short-term memory: A reconsideration of mental storage capacity. Behavioral and brain sciences, 24(1), 87–114.

20. Knill, D. C., & Pouget, A. (2004). The Bayesian brain: the role of uncertainty in neural coding and computation. TRENDS in Neurosciences, 27(12), 712–719.

21. Körding, K. (2007). Decision theory: what” should” the nervous system do? Science, 318(5850), 606–610.

22. Schurgin, M. W., Wixted, J. T., & Brady, T. F. (2020). Psychophysical scaling reveals a unified theory of visual memory strength. Nature Human Behaviour, 1–17.

23. Vul, E., Hanus, D., & Kanwisher, N. (2009). Attention as inference: selection is probabilistic; responses are all-or-none samples. Journal of Experimental Psychology: General, 138(4), 546.

24. Vul, E., & Pashler, H. (2008). Measuring the crowd within: Probabilistic representations within individuals. Psychological Science, 19(7), 645–647.

25. Bays, P. M. (2014). Noise in neural populations accounts for errors in working memory. Journal of Neuroscience, 34(10), 3632–3645.

26. Bays, P. M. (2015). Spikes not slots: noise in neural populations limits working memory. Trends in cognitive sciences, 19(8), 431–438.

27. Honig, M., Ma, W. J., & Fougnie, D. (2020). Humans incorporate trial-to-trial working memory uncertainty into rewarded decisions. Proceedings of the National Academy of Sciences, 117(15), 8391–8397.

28. Keshvari, S., Van den Berg, R., & Ma, W. J. (2012). Probabilistic computation in human perception under variability in encoding precision. PLoS One, 7(6).

29. Rademaker, R. L., Tredway, C. H., & Tong, F. (2012). Introspective judgments predict the precision and likelihood of successful maintenance of visual working memory. Journal of Vision, 12(13), 21–21.

30. Suchow, J. W., Fougnie, D., & Alvarez, G. A. (2017). Looking inward and back: Real-time monitoring of visual working memories. Journal of Experimental Psychology: Learning, Memory, and Cognition, 43(4), 660.

31. Wan, X., Wang, W., Liu, J., & Tong, T. (2014). Estimating the sample mean and standard deviation from the sample size, median, range and/or interquartile range. BMC medical research methodology, 14(1), 135.

32. Massey Jr, F. J. (1951). The Kolmogorov-Smirnov test for goodness of fit. Journal of the American statistical Association, 46(253), 68–78.

33. Suchow, J. W., Brady, T. F., Fougnie, D., & Alvarez, G. A. (2013). Modeling visual working memory with the MemToolbox. Journal of Vision, 13(10), 9–9.

34. Schneegans, S., & Bays P. M. (2016). No fixed item limit in visuospatial working memory. Cortex, 83, 181–193.

35. Oberauer, K., & Lin, H. Y. (2017). An interference model of visual working memory. Psychological Review, 124(1), 21.

36. Pratte, M. S. (2019). Swap errors in spatial working memory are guesses. Psychological Bulletin & Review, 26(886), 958–966.

37. Matthey, L., Bays, P. M., & Dayan, P. (2015). A probabilistic palimpsest model of visual shortterm memory. PLoS Computational Biology, 11(1).

38. Swan, G., & Wyble, B. (2014). The binding pool: A model of shared neural resources for distinct items in visual working memory. Attention, Perception, & Psychophysics, 76(7), 2136–2157.

39. Sutterer, D. W., Foster, J. J., Adam, K. C., Vogel, E. K., & Awh, E. (2019). Item-specific delay activity demonstrates concurrent storage of multiple active neural representations in working memory. PLoS Biology, 17(4), e3000239.

40. Johnson, J. S., Spencer, J. P., & Schöner, G. (2006). A dynamic neural field theory of multi-item visual working memory and change detection. In Proceedings of the Annual Meeting of the Cognitive Science Society (Vol. 28, No. 28).

41. Serences, J. T. (2016). Neural mechanisms of information storage in visual short-term memory. Vision research, 128, 53–67.

42. Sreenivasan, K. K., & D’Esposito, M. (2019). The what, where and how of delay activity. Nature Reviews Neuroscience, 20(8), 466–481.

